# Environmental drivers of genetic adaptation in two corals from the Florida Keys

**DOI:** 10.1101/2025.03.03.641296

**Authors:** Kristina L. Black, J.P. Rippe, Mikhail V. Matz

**Affiliations:** Department of Integrative Biology, University of Texas at Austin, Austin, TX, USA

**Keywords:** Seascape genomics, 2bRAD, random forest, cryptic speciation, polygenic adaptation

## Abstract

Increasingly frequent marine heatwaves devastate coral reefs around the world, so there is great interest in finding warm-adapted coral populations that could be used as sources for assisted gene flow and restoration. Here, we evaluated the relative power of various environmental factors to explain coral genetic variation, suggestive of differential local adaptation to these factors, across the Florida Keys Reef Tract. We applied a machine learning population genomic method (RDAforest) to two coral species - the mustard hill coral *Porites astreoides* and the lettuce coral *Agaricia agaricites* - sampled from 65 sites covering the whole reef tract. Both species comprised three genetically distinct lineages distributed across depths in a remarkably similar way. Within these lineages, there was additional genetic variation explained by depth, but even more within-lineage variation was cumulatively explained by water chemistry parameters related to nitrogen, phosphorus, silicate, and salinity. Visualizing the predicted environment-associated genetic variation on a geographic map suggests that these associations reflect adaptation to certain aspects of the inshore-offshore environmental gradient, and, to a lesser extent, to difference of Middle and Lower Keys from the rest of the reef tract. Thermal parameters, most notably maximal monthly thermal anomaly, were also consistently identified as putative drivers of genetic divergence, but had a relatively low explanatory power compared to depth and water chemistry. Overall, our results indicate that temperature is not the most important driver of differential coral adaptation in the Florida Keys, and underscore depth and water chemistry as more important environmental factors from the corals’ perspective. These findings are relevant for planning assisted gene flow and restoration efforts: not just any warmer reef would be an optimal source of transplants to promote thermal adaptation within the local coral population.

## Introduction

Coral reefs are rapidly declining worldwide (Hughes et al. 2017; Sully et al. 2019), but there is growing interest to understand how scleractinian corals could adapt to environmental stressors such as climate change (Funk et al. 2019; Munday et al. 2013). Coral restoration efforts aim to facilitate adaptation by assisting gene flow between populations with variable heat tolerance (Aitken and Whitlock 2013; Caruso, Hughes, and Drury 2021; van Oppen et al. 2017). However, in a heterogeneous seascape composed of complex gradients of environmental differences, it is firstly important to determine which environmental factors actually drive genetic adaptation.

Thus far, studies of coral adaptation have been almost entirely focused on thermal parameters, since rising sea surface temperature is perceived as the foremost threat to corals under climate change (van Woesik, Shlesinger, and Grottoli 2022). Such studies typically compare corals from a few (often, just two) locations differing in mean temperature (Howells et al. 2016; Dixon et al. 2015), temperature variability (Barshis et al. 2013; Bay and Palumbi 2014; Carly D. Kenkel and Matz 2016; Thomas et al. 2018), and/or frequency of temperature extremes (Thomas et al. 2022). While such experimental design can test whether the targeted gradient aligns with genetic adaptation, it cannot ascertain that the adaptation is indeed driven by the targeted parameter, as opposed to some other environmental difference between chosen locations. To disentangle effects of different parameters on coral adaptation, the study must be designed quite differently, following the landscape/seascape genomics paradigm (Selmoni, Vajana, et al. 2020): instead of deep sampling of a few locations representing the hypothetically important gradient, the design must maximize the number of locations sampled to cover the whole diversity of environments in the region, taking only a few or even just one sample per location. With such experimental design it is then possible to identify which ones of the many (often correlated) environmental gradients are the best predictor of genetic divergence. With this knowledge, it is then possible to predict where in the seascape divergent adaptation is expected and, using future values for the environmental parameters, where populations will be the most threatened by climate change (Fitzpatrick and Keller 2015). The power of this approach is best illustrated by the paper by Bay et al. (2018), where the authors identified adaptation gradients and predicted population declines in yellow warbler (a migratory songbird), which were confirmed by on-the-ground observations.

Seascape genomics studies of corals were pioneered by Oliver Selmoni and Stephane Joost (Selmoni, Rochat, et al. 2020; Selmoni et al. 2021) and still are few; moreover, they tend to be focused on thermal parameters. To date, the general picture of what actually drives differential coral adaptation in the wild has not yet emerged, and every new study contributes a new piece to this puzzle. One notable pattern that has been observed in multiple studies is the role of depth in driving genetic divergence in a variety of coral species across the globe (Bongaerts et al. 2017; Prada and Hellberg 2021; Rippe et al. 2021; Bongaerts et al. 2011; Johnston et al. 2022; Carlon and Budd 2002; Cabacungan et al. 2025). Here, we used a recently developed machine-learning-based seascape genomics approach, RDAforest (Matz and Black 2024), to identify drivers of local adaptation in two species of corals from the Florida Keys Reef Tract (FKRT), *Agaricia agaricites* and *Porites astreoides*, sampled from 65 sites spanning the entire extent of the FKRT (Fig. 1 A)

**Figure 1:**
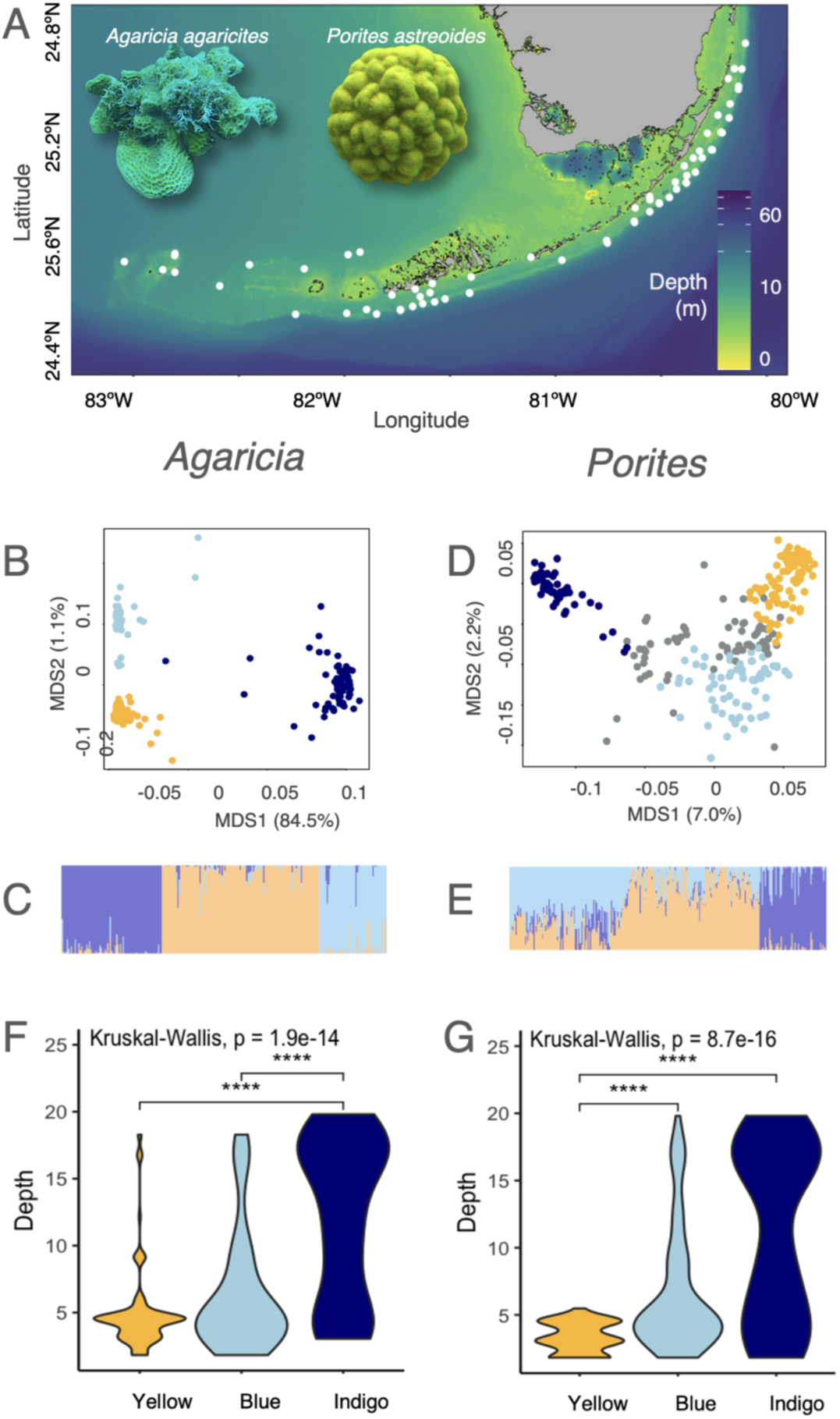
Sampling locations and genetic structure associated with depth. (A) Two ubiquitous coral species-*Agaricia agaricites* and *Porites astreoides* - were sampled across 65 sites spanning the entire Florida Keys Reef Tract. (B, C) Genetic structure of *A. agaricites* (n=250). (D, E) Genetic structure of *P. astreoides* (n=269). (B, D) Principal coordinate analyses based on the identity-by-state genetic distance matrices, colored by admixture assignment, with individuals with <50% assignment to any lineage shown in grey. (C, E) Admixture barplots showing assignment of each individual to the three lineages. (F, H) Violin plots showing the range of depths inhabited by each admixture cluster. For both species, the shallow specialist cluster is depicted in yellow, the shallow-preferring cluster is blue, and the depth-tolerant cluster is indigo.

FKRT is one of the world’s largest barrier reefs in the world, which experienced a drastic decline in coral cover in recent years (Jones et al. 2022; Lauren T. Toth et al. 2022). The FKRT has a long history of temperature stress, as reef accretion began declining in the Holocene due to climatic cooling (Lauren T. Toth et al. 2018). Due to its subtropical location, the reef often experiences cold anomalies that lead to coral mortality (Kemp et al. 2016), while modern climate change brings increasingly frequent heat waves causing severe bleaching events (Manzello 2015). Restoration is the primary approach for enhancing coral populations in the FKRT (Ladd, Burkepile, and Shantz 2019), and it has the potential to significantly increase reef-accretion (Lauren T. Toth et al. 2022). Molecular tools are becoming increasingly useful for identifying patterns of local adaptation and selecting coral genotypes for breeding and outplanting (Baums 2008). But survival of outplanted corals varies across the FKRT (Drury and Lirman 2021), even at sites that are adjacent to each other (Banister and van Woesik 2021), so there is a need to strategize the spatial arrangement of outplants to ensure the highest survival. Indeed, restoration efforts could be optimized by either maximizing the genetic diversity outplanted across the entire reef, or by targeting outplanting efforts to place corals where they would be genetically pre-adapted to the local conditions (Baums et al. 2019).

Researching coral genotype-environment associations across the Florida seascape poses challenges as most of the local coral species have become scarce. Major reef-builders of the genus *Acropora* (Cramer et al. 2020) and other iconic species (Neely et al. 2021) have declined by at least 55% in reef assemblages in recent decades (L. T. Toth et al. 2019). Nonetheless, *Agaricia agaricites* and *Porites astreoides* are still relatively ubiquitous. Both are brooding species (i.e., they reproduce by releasing fully-formed larvae that already contain algal symbionts) that are relatively stress tolerant and were even labeled “weedy” for their persistence despite climate change, disease, and continuous anthropogenic disturbances (L. T. Toth et al. 2019). *P. astreoides*, in particular, increased by at least 29% in the relative composition of coral assemblages (L. T. Toth et al. 2019). These resilient species may hold valuable insights to the patterns of local adaptation and which environmental gradients drive genetic differentiation across the reef tract.

## Materials and methods

### Sample collection and DNA sequencing

In June 2019, samples of four to five colonies of *A. agaricites* and *P. astreoides* were collected from 65 sites spanning the entire FKRT (Fig. 1 A). Samples were preserved in 100% ethanol and stored at -80C. Genomic DNA was isolated using a CTAB procedure (Baker and Cunning 2016) for 2bRAD sequencing (S. Wang et al. 2012). 2bRAD is a reduced-representation sequencing approach of the family of restriction site associated DNA (RAD) methods, designed to reproducibly sequence approximately 0.5% of the genome. 2bRAD libraries were prepared according to the updated protocol (hosted at https://github.com/z0on/2bRAD_denovo), allowing for in-read barcoding and removal of PCR duplicates. The libraries were sequenced on the Illumina NovaSeq SR100 platform at the Genomic and Sequencing Analysis Facility at the University of Texas at Austin. Reads were trimmed and demultiplexed, then compiled into a cluster-derived reference, or CDR (S. Wang et al. 2012), following the *de novo* pipeline https://github.com/z0on/2bRAD_denovo. The CDR was concatenated with genomes of the four major clades of zooxanthellae symbionts for mapping (NCBI accession for *Symbiodinium* reference: GCA_003297005.1; *Brevolium*: GCA_000507305.1; *Cladocopium*: GCA_003297045.1; *Durisdinium*: GAFP00000000). Reads were mapped to this reference with bowtie2 (Langmead and Salzberg 2012). On average, 1,151,112 reads per sample of *A. agaricites* and 1,507,436 reads of *P. astreoides* aligned to their respective coral CDR. Reads that mapped to symbiont genomes (*A. agaricites*: mean=1,467, *P. astreoides*: mean=1,537) were removed. To filter samples by sequencing depth, individuals with less than a quarter of sites at 5x coverage were discarded. This retained 253 out of 289 samples of *A. agaricites*, and 299 out of 330 samples of *P. astreoides*. We used ANGSD (Korneliussen, Albrechtsen, and Nielsen 2014) to remove low quality SNPs (with >0.1% chance of sequencing error and >0.1% chance of erroneous mapping), sites not genotyped in at least 75% of individuals (to remove non-coral contaminants), and sites with a minor allele frequency < 0.05. After filtering, we derived identity-by-state (IBS) genetic distances between samples and evaluated relatedness among samples using ngsRelate (Korneliussen and Moltke 2015).

### Population Structure

The shape of the data captured by the first two principal coordinates (Fig. 1 C, E) strongly suggested the presence of at least three distinct genetic groups within each species. To confirm this more formally, we explored admixture scenarios k=2-3 using NGSadmix (Skotte, Korneliussen, and Albrechtsen 2013). We also followed the cluster analysis pipeline from (Borcard, Gillet, and Legendre 2011). We assessed four clustering methods: single linkage (a.k.a. nearest neighbor sorting, where individuals are clustered based on shortest pairwise distance), complete linkage (a.k.a. furthest neighbor sorting, where individuals are clustered based on maximum distance), UPGMA (where individuals are clustered based on mean distance), and Ward clustering (based on least squares). For each clustering method, we computed the cophenetic matrix (a matrix of distances based on the inferred tree structure), and chose the optimal method (UPGMA) by the highest correlation between the cophenetic matrix and the original distance matrix. We then determined the optimal number of clusters based on two criteria. The first one was the widest silhouette width, a value between -1 to 1 that reflects the difference between within-cluster distances and distances to members of the next closest cluster (Borcard, Gillet, and Legendre 2011). The second criterion was the highest Mantel correlation between original genetic distance matrix and binary dissimilarity matrix computed to represent cluster membership (i.e. distances to members of the object’s own cluster were set to 0, the rest were set to 1).

We examined the initial hierarchical clustering based on IBS and relatedness to remove highly similar samples that could disrupt ordinations. *P.* astreoides contained many highly genetically related individuals, with 23 clonal groups and 19 potential full and half sibling groups that shared 25-50% of alleles (Supplementary Fig. 1). By contrast, we observed only two clonal groups and seven potential sibling groups in *A. agaricites* (Supplementary Fig. 1). In both species, only one representative with the highest sequencing depth from each clonal group was retained for genotype-environment analyses. To remove highly related individuals, we visually inspected a dendrogram of hierarchical clustering and pruned branches with >0.8 similarity. Final sample sets for each species and genetic lineage were verified in a principal coordinate analysis (PCoA) so that remaining putatively-related groups did not distort population structure. After filtering, we retained 11,332 SNPs across 250 individuals for *A. agaricites* and 5,332 SNPs across 269 individuals for *P. astreoides* for producing the final IBS distance matrices for each species.

To isolate specific genetic lineages for genotype-environment association analysis, we first identified a subset of individuals with high admixture assignment to the lineage and then remade IBS distance matrices for each. We evaluated the hierarchical clustering based on per-lineage IBS and removed potential related individuals (if detected) before performing genotype-environment association analyses on each lineage independently.

Isolation-by-distance within each species and lineage was assessed visually by plotting genetic distances against geographic distances. Significance of association between genetic and geographic distances was formally evaluated using Procrustes test (*vegan:protest* version 2.5.7, (Oksanen et al. 2007)) on two distance-based ordinations: genetic (IBS genetic distance) and spatial (latitude and longitude). Since significant trends have been detected in all cases, we have used latitude and longitude (converted to universal transverse mercator projection) as covariates while constructing ordinations for the genotype-environment association analyses.

### Environmental variables

#### Depth

For the 65 sampling sites, depth was recorded *in situ*. For the FKRT seascape, the depth raster with 0.001° geospatial resolution was obtained from the General Bathymetric Chart of the Oceans website (https://www.gebco.net/).

#### *In situ* water quality data

Water chemistry, bottom temperature, and water column stratification data for 224 sites positioned throughout the Florida Keys were obtained from SERC Water Quality Monitoring Network (http://serc.fiu.edu/wqmnetwork/, (Boyer 2003). Surface measurements were excluded if bottom measurements for the same parameter were available. We split the data into two time slices, 1995-2007 (“past”) and 2008-2021 (“present”). For each parameter, we calculated the median and 0.1-0.9 interquantile range at each station over the whole time slice. To project the data from discre<u>t</u>e SERC stations onto the whole FKRT seascape and identify values of each parameter at our sampling sites, we have used kriging interpolation with regularization (https://github.com/z0on/smoothKrige). The original SERC data were first smoothed with a 5 km gaussian kernel (function *spatstat::Smooth*), which corresponds to the 5km resolution of the target FKRT raster, and then interpolated using *automap::autokrige* function (Hiemstra et al. 2009). This two-step procedure was adopted because Interpolation without smoothing occasionally produced nonsensical results, based on visualizing the resulting rasters (see examples at https://github.com/z0on/smoothKrige).

#### Satellite-based data

Daily sea surface temperature (sst), thermal anomaly (dhw), diffuse attenuation coefficient at 490nm (k490), and <u>c</u>hlorophyll A concentration data were obtained from NOAA’s ERDDAP server (Simons 2017) using *rerddap::ed_search_adv* function. The thermal dataset was NOAA_DHW_monthly, the k490 dataset was erdMH1kd4901day, and the chlorophyll A dataset was erdMH1chlamday. We used two five-year time slices for purposes of comparison, 01/01/2003-12/31/2007 (“past”) and 01/01/2015-12/31/2019 (“present”). For each pixel of the sst raster we calculated minimum and maximum monthly means for each year and then averaged these values across years. For thermal anomaly, we calculated the maximum monthly mean (dhw_max) for each year and then averaged across years. k490 and chlorophyll <u>a</u> were summarized as yearly medians and 0.1-0.9 interquantile ranges, which were averaged over years. Chlorophyll A data were then dropped because they were strongly correlated with k490 data (Pearson’s *r* > 0.9).

#### Reducing multicollinearity

*In situ* and satellite data were merged to produce rasters covering the whole FKRT with 0.009° geospatial resolution. We then examined all the collected variables for multicollinearity, by performing a hierarchical clustering based on absolute value of Pearson correlation (function *stats::hclust*, method= “complete”) and examining the tree (Supplementary Fig. 2). Several groups of parameters were identified with absolute correlation of 0.9 or higher; we have chosen just one representative variable for each such group. As a result, we ended up with 31 parameters to predict coral adaptation (Table 1, medians and 0.1-0.9 interquantile ranges for all except depth, sst, and dhw_max).

**Table 1:** Environmental variables used as predictors of coral genetic structure.

| ID | Type | Environmental variable |
| --- | --- | --- |
| sst | Therm | *Sea surface temperature (satellite)*<br>min : *minimal monthly mean*<br>max : *maximal monthly mean* |
| dhw_max | Therm | *maximum monthly thermal anomaly* |
| k490 | Phys | *Diffuse attenuation coefficient at 490nm, proxy of turbidity.*<br>*Highly correlated with Chlorophyll A.* |
| depth | Depth | *Depth* |
| NO3 | Chem | *Nitrates* |
| NO2 | Chem | *Nitrites* |
| NH4 | Chem | *Ammonium* |
| TN | Chem | *Total nitrogen* |
| DIN | Chem | *Dissolved inorganic nitrogen* |
| TP | Chem | *Total phosphorus* |
| DIN.TP | Chem | *Dissolved inorganic nitrogen : total phosphorus ratio. Estimate of nutrient limitation of phytoplankton* |
| SRP | Chem | *Soluble reactive phosphorus* |
| SiO2 | Chem | *Silicate* |
| SAL | Chem | *Salinity* |
| TEMP | Therm | *in situ bottom temperature* |
| NP | Chem | *Nitrogen : phosphorus ratio* |
| TN.TP | Chem | *Total nitrogen : total phosphorus ratio. Proxy of nutrient limitation of phytoplankton* |
| Si.DIN | Chem | *Silicate : nitrogen ratio* |
| DISGT | Phys | *Delta sigma-t: measure of water column stratification* |

### Genotype-environment associations

To identify the most important environmental predictors of genetic divergence and place putatively adaptive genetic variation on a geographic map, we used RDAforest (Matz and Black 2024), https://github.com/z0on/RDA-forest), which predicts scores along principal axes of genetic variation (“genetic PCs”, or gPCs) using random forest regressions against multiple environmental parameters. Using random forest instead of linear regressions gives RDAforest the ability to detect linear, non-linear, and non-monotonous associations, while at the same time accounting for all possible interactions between predictors. By focusing on overall genetic structure rather than individual SNP-environment associations, RDAforest capitalizes on the predominantly polygenic nature of adaptation (Berg and Coop 2014). This is fundamentally similar to generalized dissimilarity modeling (Ferrier et al. 2007) and the application of classical redundancy analysis (Borcard, Gillet, and Legendre 2011) to the genotype-environment association problem (Capblancq and Forester 2021). In addition, RDAforest uses jackknifing to model the uncertainty of reconstructing gPCs while regressing out the effects of nuisance variables (such as spatial coordinates). We accounted for isolation-by-distance using linear regression to remove the effect of geographic coordinates (Matz and Black 2024). Besides spatial coordinates, additional covariates that were regressed out included posterior probabilities of being assigned to other cryptic lineages. RDAforest also incorporates a procedure to identify likely truly influential predictors among multiple correlated ones based on their response to the *mtry* parameter setting (Matz and Black 2024).

### Mapping adaptive neighborhoods

After identifying the influential predictors based on observed data at the sampled sites, RDAforest can predict gPC scores at unsampled locations based on environmental parameters there. Such predictions can then be visualized on a geographic map, which we call a “map of adaptive neighborhoods”, with contrasting colors signifying genetic divergence (Matz and Black 2024).

Here we must note that RDAforest does not extrapolate. When encountering a predictor value outside the predictor’s range observed during model construction (i.e., across sites sampled for genetic analysis), the default behavior of RDAforest is to drop the location where this occurs. This conservative behavior may result in too many missing spatial points to form an informative map, especially when the RDAforest model includes many predictors. To deal with this problem, we extended the allowed predicted range by 10% of the sampled range in both directions. For prediction purposes, predictor values falling in these 10% margins were treated as either minimal or maximal observed values (depending on the direction of their deviation from the sampled range).

To emphasize the most important features of the map, RDAforest clusters spatial points which have similar values for predicted gPCs. Here, we clustered spatial points based on turnover curves (Ellis, Smith, and Pitcher 2012), followed by merging the clusters based on random forest predictions. This two-step procedure tends to generate less noisy clusters than direct clustering of random forest predictions (Matz and Black 2024).

### Genetic offset

Seascape-wide predictions can be computed based on environmental data from different times. The Euclidean distance between gPC predictions for different times is “genetic offset” (Fitzpatrick and Keller 2015), which illustrates how much genetic adaptation will be (or was) necessary to adapt through time at a given location. The map of genetic offset is sometimes called a “genomic vulnerability” map (Bay et al. 2018), because it shows the risk of maladaptation as time goes on. Here, we computed genetic offsets between pre-2008 and post-2008 seascapes.

### Environmental mismatch

Environmental mismatch, computed by the function *RDAforest::env_mismatch*, is the Euclidean distance between a given gPCs vector (corresponding to the genotype of a specific individual or predicted for a location it was sampled from) and gPCs predictions for the rest of the modeled range (Matz and Black 2024). A map of environmental mismatch illustrates where the individual in question is predicted to be more or less maladapted if transplanted.

## Results

### Cryptic genetic structure

In *Agaricia*, three clusters were visually apparent on the plot of the first two principal coordinates of the identity-by-state distance matrix (Fig. 1 B). The analogous plot for *Porites* looked like a V-shaped cline (Fig. 1 D), suggesting existence of two genetic lineages (tips of the “V”) that were not interbreeding with each other but both interbred with a third lineage (the base of the “V”). To determine the optimal number of clusters more formally, we first chose the optimal clustering method, the one producing the highest cophenetic correlation with original genetic distances. UPGMA (*A. agaricites* R=0.990, *P. astreoides* R=0.872) outperformed single linkage (*A. agaricites* R=0.983, *P. astreoides* R=0.571), complete linkage (*A. agaricites* R=0.988, P*. astreoides* R=0.692), and Ward clustering (*A. agaricites* R=0.983, *P. astreoides* R=0.705) in both species. We then used UPGMA to determine the optimal number of clusters (*k*) by the widest silhouette width. After testing all possible values of *k* the widest silhouettes were observed at *k*=3 for both species (Supplementary Fig. 3). Three clusters also produced the highest Mantel correlation between the binary matrix of cluster memberships and the original matrix of genetic distances (*Agarcia*: R=0.92, *Porites*: R=0.72). We therefore assumed that we have three genetic lineages in both species, and assigned individuals to them based on ngsAdmix results for k=3 (Fig. 1 C, E).

In both species, the three genetic lineages were largely sympatric across sampled sites (Supplementary Fig. 4) and had remarkably similar depth distribution (Fig. 1 F, G). One of the lineages, designated “yellow” in both species, was rarely (if ever) found deeper than 5m. In contrast, a lineage designated “indigo” was predominantly found at depths exceeding 15m. The third lineage, designated “blue”, was preferentially found in shallow waters but also across the whole depth range. In both species, indigo was the most genetically distinct lineage (Fig. 1 B, D).

### Genotype-environment associations

We ran RDAforest on the full sample sets for each species to identify the major environmental predictors of between-lineage divergence. We then also ran the analysis separately for the individual cryptic lineages, to reveal putative adaptive divergence within them. For *A. agaricites,* yellow and indigo lineages were represented by individuals with >90% probability assignment based on ngsAdmix results (yellow: n=92, indigo: n=60). For the blue lineage of *A. agaricites*, to boost sample size, we used samples with >60% assignment (n=48). For *P. astreoides*, we individually analyzed the indigo lineage (>60% assignment, n=53) and blue+yellow lineages together since they did not show clear separation (Fig. 1 D; <50% assignment to indigo, n=192). Complete data per species as well as all by-lineage subsets showed significant isolation-by-distance (Procrustes test for correlation between genetic and geographic distances, p<0.05); to account for that, we regressed spatial coordinates out during construction of genetic ordination.

In addition, ngsAdmix-generated probabilities of assignment to other cryptic lineages were used as covariates when analyzing individual lineages.

#### Overall model fit

The proportion of total variation explained by RDAforest models using predictors retained after *mtry*-based selection procedure ranged from 0.33 (*A. agaricites* full dataset) to just 0.01 (*P. astreoides* indigo). *A. agaricites* datasets generated considerably better model fit compared to *P. astreoides*, with almost 10-fold larger proportion of variation explained (Fig. 2).

**Figure 2.**
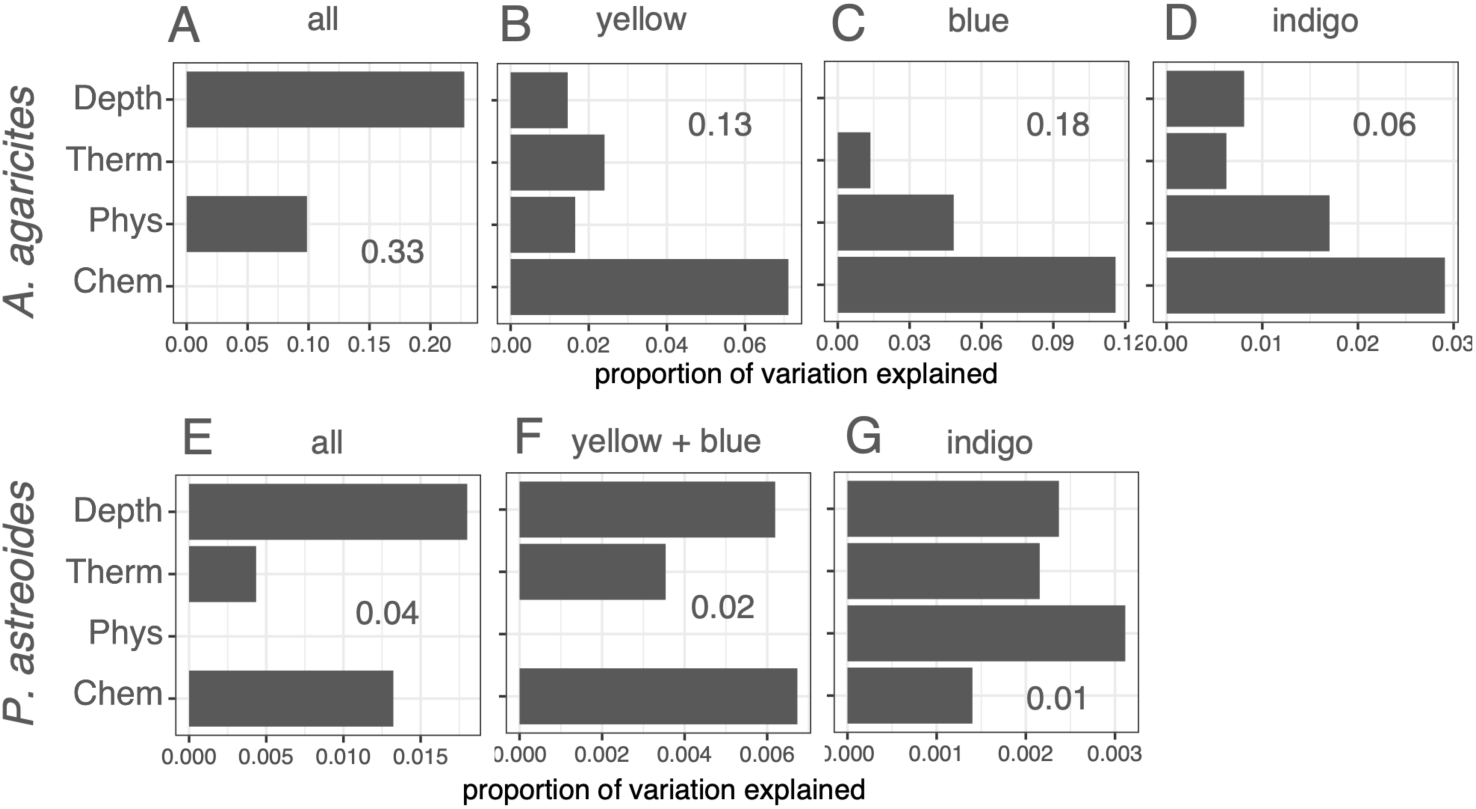
Proportions of genetic variation explained by RDAforest models, summed up by the type of predictor. The predictor types (y-axis categories) are Depth, thermal parameters (“Therm”: minimum monthly temperature, maximum monthly temperature, maximum monthly thermal anomaly), other physical parameters (“Phys”: turbidity, water column stratification), and water chemistry (“Chem”: remaining predictors related to N, P, Si and their ratios). The analysis either used complete datasets for the coral species (A, E) or subsets corresponding to cryptic lineages, indicated above the plots (B-D, F, G). (A - D) *A. agaricites*, (E - G) *P. astreoides*. The value inside each plot is the total proportion of variation explained by the RDAforest model using predictors passing *mtry*-based selection procedure.

#### Depth

For both species, depth was by far the most important explanatory parameter for full datasets, explaining 2.5-3-fold more variation than the next best predictor (Fig. 2 A, E, Supplementary Fig. 2). Depth was also the far-leading predictor for *P. astreoides* blue+yellow subset, reflecting segregation of blue and yellow lineages by depth (Fig. 1 G). Notably, depth was also an important predictor within individual lineages, with the only exception of *A. agaricites* blue (Fig. 2, Supplementary Fig. 5). Turnover curves for depth (Fig. 3) reveal at which depths major genetic transitions happen: such transitions appear as particularly large steps in the curve. For the complete *A. agaricites* set, the major threshold was at 11m (Fig 3 A), corresponding to the divergence of indigo lineage from the other two (Fig. 1 F). The same 11m threshold is also visible within *A. agaricites* indigo lineage (Fig. 3 C), while yellow lineage shows a threshold at 8m (Fig. 3 B). A similar 7-8m threshold is also characteristic of the *P. astreoides* yellow+blue subset (Fig. 3 E). *P. astreoides* indigo lineage shows the shallowest threshold, between 4 and 5m (Fig. 3 F). No obvious depth thresholds are discernible in the complete *P. astreoides* dataset (Fig. 3 D).

**Figure 3:**
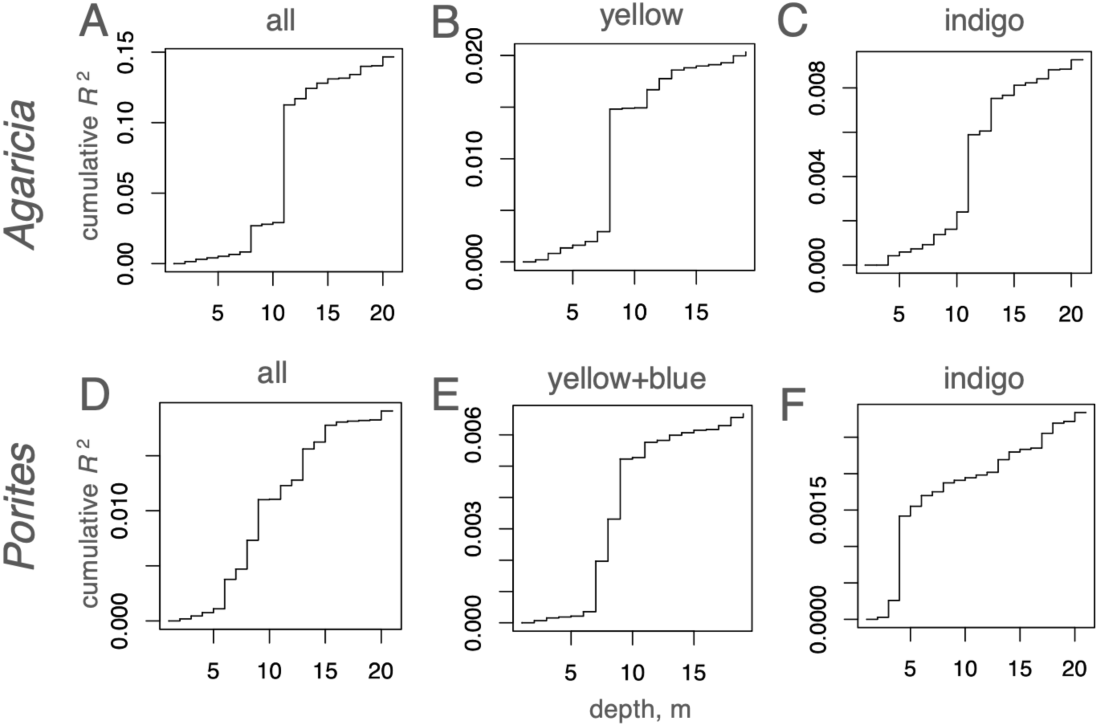
Depth turnover curves (cumulative genetic variation captured as one moves across the predictor range). (A-C) *Agaricia*, (D-F) *Porites*. Label on top of each panel identifies the sample set: either whole species (A, D) or cryptic lineages (B, C, E, F).

#### Temperature

Thermal parameters were selected as important predictors in all analyses except the full *A. agaricites* dataset. Maximum monthly thermal anomaly (dhw_max) was selected most often, in all three *A. agaricites* lineages and also in full and yellow+blue datasets of *P. astreoides*. The turnover curve for this parameter for *A. agaricites* yellow lineage demonstrated a notable transition between 1.45-1.50 °C; turnover curves for other lineages were smooth, lacking major steps (Supplementary Fig. 6). The amount of variation explained by thermal parameters was typically the lowest of the four predictor types considered, with the exception of *A. agaricites* yellow (second most important) and *P. astreoides* indigo (third most important).

#### Water chemistry

Predictors related to nitrogen, phosphorus, silicate, salinity, and their ratios dominated models for individual *A. agaricites* lineages (Fig. 2 B-D) and were also important (on par with depth) for *P. astreoides* whole dataset and yellow+blue dataset (Fig. 2 E,F).

#### Physical parameters other than temperature

These included diffuse attenuation coefficient at 490 nm (k490), which is a proxy or turbidity, and delta sigma-t (DSIGT), a measure of water column stratification. These parameters explained moderate amounts of variation in *A. agaricities* lineages and *P. astreoides* indigo. Still, the yearly range of turbidity (k490_range) was the only retained predictor in addition to depth in the model for the full *A. agaricites* dataset, explaining 10% of total variation (Fig. 2 A and Supplementary Fig. 2 A).

#### Maps of adaptive neighborhoods

Geographic maps colored according to the first three principal axes of predicted gPCs (“adaptive neighborhoods” maps) generally followed the inshore-offshore gradient, with differences accumulating as one moves seaward from the main island chain. The only exception was *A. agaricites* indigo (Fig. 4 D), which instead emphasized the distinction between the middle region of the island chain (Middle and Lower Keys) and the rest of the environment. The same differentiation (Middle and Lower Keys vs. the rest) was also visible for *A. agaricites* yellow (Fig. 4 B) and *P. astreoides* indigo (Fig. 4 G; for this lineage many Middle and Lower Keys areas were not predictable because predictor values there were outside the range observed for field-collected samples).

**Figure 4:**
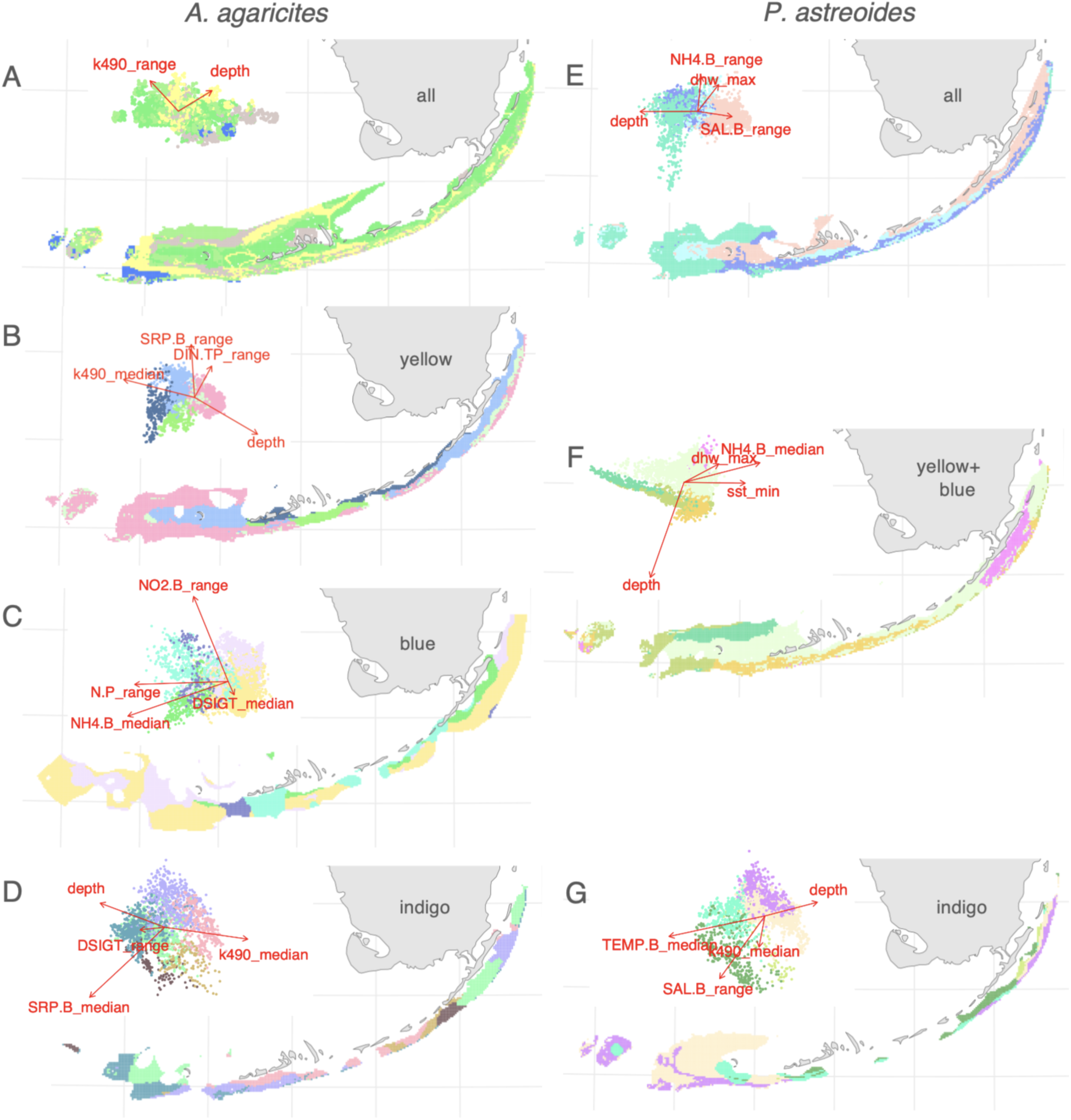
Maps of environment-associated genetic variation (“adaptive neighborhoods”). (A-D) *Agaricia*, (E-G) *Porites.* Colors on the maps correspond to predicted scores along the first three principal axes of environment-associated genetic variation (insets show the first two axes). Contrasting colors in each PCA (upper left of each panel) suggest differential adaptation, putatively driven by the corresponding environmental vectors. Those same colors plotted on the map indicate regions where corals may be locally adapted to their environmental predictors. Predictions are made only for points where the predictor values do not fall outside their range across sampled sites (Fig. 1 A) plus-minus 10% margin. Arrows on the insets are directions of the most important environmental gradients, aligned with the predicted principal axes via linear regression. Sample set is identified by the black text label inside the Florida contour: panels A and E show predictions based on all samples, other panels show predictions based on subsets of samples corresponding to cryptic lineages (Fig.1 C, E).

#### Distance to land

To confirm the importance of specific environmental parameters rather than just the distance to the main Florida Keys island chain (the simplest proxy of the inshore-offshore gradient), we reran all the analyses including “distance to land” (dis2land) as one of the predictors. This predictor was the distance from sampling location to the narrow continuous polygon encompassing the island chain from Key Biscayne to Marquesas Keys, with a separate polygon for Dry Tortugas. Including dis2land did not increase the proportion of variation explained by any of our models. In three cases (*A. agaricites* full dataset, *A. agaricites* blue, and *P. astreoides* yellow+blue) dis2land did not pass the *mtry* selection criteria. In analyses retaining dis2land among important predictors it played a minor role, not displacing the predictors chosen originally (Supplementary Fig. 7). The only analysis where dis2land became the most important predictor was *P. astreoides* indigo, but even in this case dis2land did not displace original predictors and did not change the overall map of adaptive neighborhoods (Supplementary Fig. 8).

#### Genetic offset

We computed genetic offset, comparing predictions based on pre-2008 environmental data to predictions based on post-2008 data, for *A. agaricites* only, because the RDAforest models for that species were much more robust than for *P. astreoides*. Still, geographic maps of the offset lacked interpretable structure (Supplementary Fig. 9), suggesting that either the maladaptation risks are approximately uniform across the FKRT, or our data are not strong enough to reveal the pattern of their distribution.

#### Environmental mismatch

To give an example of practical use of our models, we generated a map of environmental suitability for a coral of *A. agaricites* indigo lineage sampled near Marquesas Keys. The map indicates that one the most suitable target areas for transplanting such a coral is in the Upper Keys, whereas many locations nearby (such as Looe Key Sanctuary Preservation Area) would be highly unsuitable (Fig. 5).

**Figure 5.**
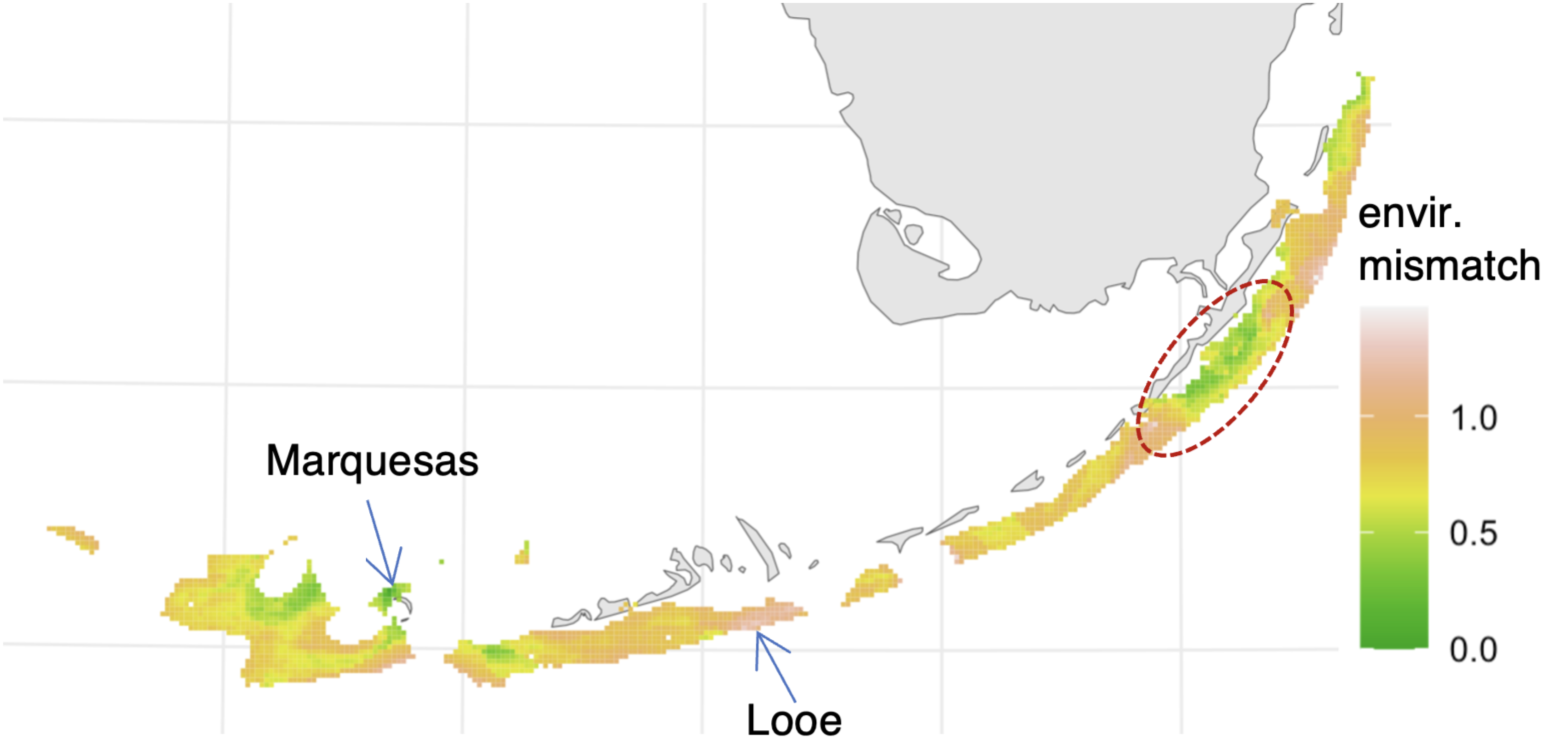
Map of environmental suitability for transplanting a coral of *A. agaricites* indigo lineage from Marquesas Keys. Darker areas are less suitable. Arrows indicate source location (Marquesas Keys) and Looe Key Sanctuary Preservation Area. Dashed outline indicates a suitable area in the Upper Keys. The environmental mismatch is scaled to the 90% quantile of the range of present-day genetic divergence predicted across the whole modeled area.

## Discussion

### Depth

We found that both *A. agaricites* and *P. astreoides* were composed of three cryptic lineages across the FKRT, with very similar partitioning of lineages across depths (Fig. 1 F, G). Indeed, depth was the strongest predictor of genetic structure in RDAforest models for both species (Fig. 2 A, D). The models also suggested that depth does not only delineate cryptic lineages; we also see differential adaptation to depth within individual lineages (Fig. 2 B, D, F). In *A. agaricites*, the depth threshold associated with genetic split between two shallow-preferring lineages and the deep-preferring one is at 11m (Fig. 3 A), while depth thresholds aligning with genetic separation within individual lineages vary from just above 5m to 11m (Fig. 3 B, C, F). Our ability to identify these thresholds illustrate the utility of turnover curves (Ellis, Smith, and Pitcher 2012) for interpreting the RDAforest models. Cryptic lineages are being discovered with increasing frequency (Struck et al. 2018), including diverse coral species (Bongaerts et al. 2017; Cooke et al. 2020; Johnston et al. 2022; Prata et al. 2022; Rippe et al. 2021; Underwood et al. 2020; Gómez-Corrales and Prada 2020; Prada and Hellberg 2013), and can be valuable sources of genetic variation for adapting to environmental change (Gómez-Corrales and Prada 2020; Rose et al. 2021; Grupstra et al. 2024). Two broadcast-spawning coral species in the Florida Keys also demonstrated genetic segregation by depth (Rippe et al. 2021). Depth is also the most important factor predicting distribution of cryptic genetic lineages of six coral species around St. Croix island (Cabacungan et al. 2025). Depth is also associated with genetic subdivision in octocorals (Prada and Hellberg 2013) and corals in the Indo-Pacific (Bongaerts et al. 2011, 2017; Carlon and Budd 2002; Gorospe, Donahue, and Karl 2015; van Oppen et al. 2018). It can be concluded that depth is consistently associated with deep splits in coral genetic structure irrespective of the coral’s life history, taxonomic position, or oceanic province. However, it remains unclear exactly which aspect of the depth gradient exerts the selective pressure driving and maintaining the genetic divergence. One possibility is that adaptation to depth is in fact adaptation to the amount and/or spectral composition of incident light, a hypothesis that can be tested in *ex situ* experiments.

### Inshore-offshore gradient

Inshore-offshore environmental gradient has long been hypothesized to be the main driver of local coral adaptation on FKRT (C. D. Kenkel, Meyer, and Matz 2013; Carly D. Kenkel and Matz 2016; Carly D. Kenkel, Almanza, and Matz 2015). Such local adaptation was documented in other coral species both on the FKRT (Rippe et al. 2021; Gallery, Rippe, and Matz 2024) and elsewhere (Underwood 2009; Matias et al. 2023; Tisthammer et al. 2021). Genetic divergence between inshore (near land) and offshore corals was visible in all our analyses except *A. agaricites* indigo (Fig. 4) and is best illustrated on the adaptive neighborhoods’ map for the whole *P. astreoides* dataset (Fig. 4 E). It is naturally associated with depth but also other variables more finely partitioning the environment across the shelf. Inshore-offshore divergence is also visible for *A. agaricites* blue lineage, which does not have depth among retained predictors (Fig. 4 C). Importantly, distance to land, when included as one of the predictors, does not absorb the predictive power of individual environmental parameters and in all cases but one (*P. astreoides* indigo) ends up playing a minor, if any, role (Supplementary Fig. 7 and Supplementary Fig. 8). This indicates that our models have likely captured some truly important environmental gradients, loosely associated with the inshore-offshore distinction.

### Middle and Lower Keys vs. the rest

The secondary pattern of environment-associated genetic differentiation that we observed was the difference of Middle and Lower Keys from the rest of the area (Fig. 4 B, D, G). Specific parameters responsible for this pattern are difficult to pinpoint in our models, but candidates include median and range of soluble reactive phosphorus (SRP.B) and median turbidity (k490_median), which could be associated with the influence of the Florida Bay. Distinction of Middle Keys from the rest of the FKRT is not surprising since this area is known to be under heavy influence of “inimical water” of the Florida Bay (Lauren T. Toth, Stathakopoulos, and Kuffner 2018), but it is unexpected that, from the corals’ perspective, Lower Keys are more similar to Middle Keys rather than to some less affected areas such as Dry Tortugas or Upper Keys.

### Water chemistry

Availability of 25 years of water chemistry measurements contributed by the Southeast Environmental Research Center’s water quality monitoring network (Boyer 2003) is a unique feature of the FKRT as the study system for coral adaptation. Water chemistry parameters together account for the largest amount of explanatory power in four out of our seven analyses (Fig. 2), but it is difficult to single out any specific ones since none of them appear to consistently rank above others in terms of predictive power (Supplementary Fig. 5). Similarly to the “why depth matters” question, this problem can be addressed in *ex situ* experiments testing influence of specific parameters on (allegedly) differentially adapted corals.

### Role of thermal parameters

The hypothesized driver of inshore-offshore genetic divergence has long been the difference in seasonal thermal variability: inshore locations experience considerably hotter summers and colder winters (Carly D. Kenkel and Matz 2016; Carly D. Kenkel, Almanza, and Matz 2015). In our RDAforest models, this variability would have been captured by the yearly range of bottom temperature, TEMP.B_range, yet we never saw it among predictors passing the *mtry*-selection criterion. A reciprocal transplantation experiment suggested that an important driver of inshore-offshore adaptation could be extremes of daily temperature range (Carly D. Kenkel, Almanza, and Matz 2015). Unfortunately, calculating this parameter requires temperature data with less than daily temporal resolution from the bottom of the water column, which were not available to us. The most commonly retained thermal parameter in our analyses was maximum monthly thermal anomaly (dhw_max), which was retained in five models out of seven (Supplementary Fig. 5). Its turnover curves were smooth except *A. agaricites* yellow, where there was an apparent transition between 1.45 and 1.50 °C (Supplementary Fig. 6). If such an anomaly was observed over the whole month, it would correspond to ∼6 degree heating weeks, a value sufficient to cause substantial bleaching (Liu and Penland 2008). Only two of our 63 sampling sites were above that threshold. Despite being the most commonly retained in the model, dhw_max explained a modest proportion of variation, typically ranking below several water chemistry parameters. Overall, our models do not provide evidence that thermal regimes are among the main drivers of local coral adaptation on the FKRT.

### Implications for coral conservation and restoration

In the context of reef conservation, the existence of genetically distinct and ecologically specialized lineages implies that nominal coral species are misleading as units of conservation (Grupstra et al. 2024). Environmental partitioning of these lineages implies that protecting a species in one type of environment might provide little benefit to representatives of the same nominal species living in another environment. In particular, the “deep refugia” hypothesis (Bongaerts et al. 2017) would not apply to our species as well as others showing strong segregation by depth (Gorospe, Donahue, and Karl 2015; Prata et al. 2022; Rippe et al. 2021; van Oppen et al. 2018); i.e. depth-tolerant corals would be unlikely to repopulate the spatially adjacent shallow-water reefs or provide genetic rescue to them, because of the “isolation by environment” (I. J. Wang and Bradburd 2014). Instead, our results suggest that corals could be moved to a location far away from their source site, as long as it represents the same or similar adaptive neighborhood. For example, deep-water *A. agaricites* (indigo lineage) from Marquesas Keys is predicted to be better adapted to the far-away Upper Keys environment rather than to the nearby Looe Key in the Lower Keys (Fig. 5). The notion that long-range mixing of populations can be achieved without disrupting adaptive patterns provides justification for advanced conservation genetics interventions, such as assisted gene flow to boost adaptive genetic diversity in managed populations (Aitken and Whitlock 2013). Our approach can inform such measures, as well as coral restoration efforts, by providing the outline of environmental gradients that matter most for the survival of outplanted corals, depending on their provenance and genetic identity. Essentially, the intersection of the cryptic lineage identity and the adaptive neighborhoods where they are found defines a genetically and adaptively uniform “stock” for coral restoration and conservation.

## Acknowledgements

The field trip for coral collection was facilitated by the International Seakeepers Society. We are deeply grateful to Capt. Jaleen Hartney who donated her personal ship’s time for this project. The project was supported by the grant from the National Science Foundation OCE-1737312 to M. V. M. The genomic analyses were performed using the high-performance computing resources of the Texas Advanced Computing Center (TACC). Environmental data were provided by NOAA’s ERDDAP server and the SERC-FIU Water Quality Monitoring Network which is supported by EPA Agreement #X994621-94-0 and NOAA Agreement #NA09NOS4260253. We are also thankful to undergraduate students who helped with DNA extractions - Sofia Beskid, Kelsey Moreland, and Ashlynn Broussard.

## Data Accessibility and benefit sharing

All sequences and metadata are deposited on the Sequence Read Archive under Bioproject PRJNA812916. All scripts, genotype data, and environment data can be found at https://github.com/z0on/FLKeys-Coral-Seascape-Gen. All contributions to this research are described in the Acknowledgements and Author contributions.

## Author contributions

M.V.M. supervised the study, and both M.V.M and J.P.R. collected the samples and provided some tools for data analysis. K.L.B. processed the samples, led the data analysis, and wrote the first version of the manuscript. All authors contributed to the final version of the manuscript.

## Supplementary Figures

**Supplementary Figure 1:**
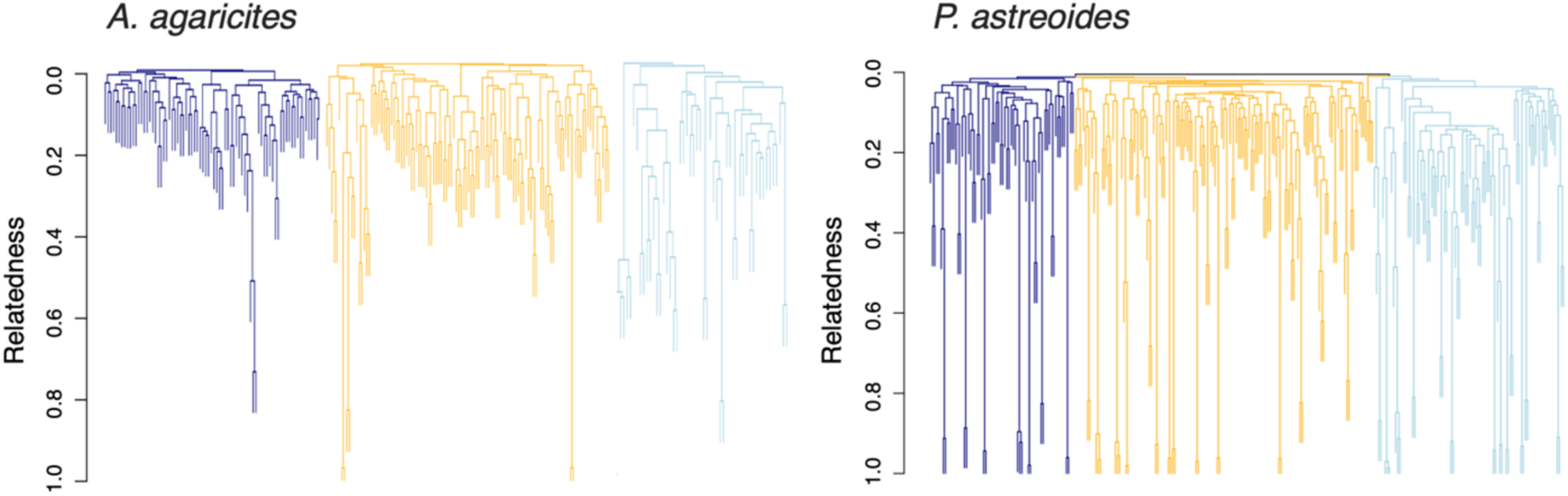
Hierarchical clustering trees based on relatedness. *P. astreoides* contains visibly more closely related and clonal (relatedness approaching 1) groups.

**Supplementary Figure 2:**
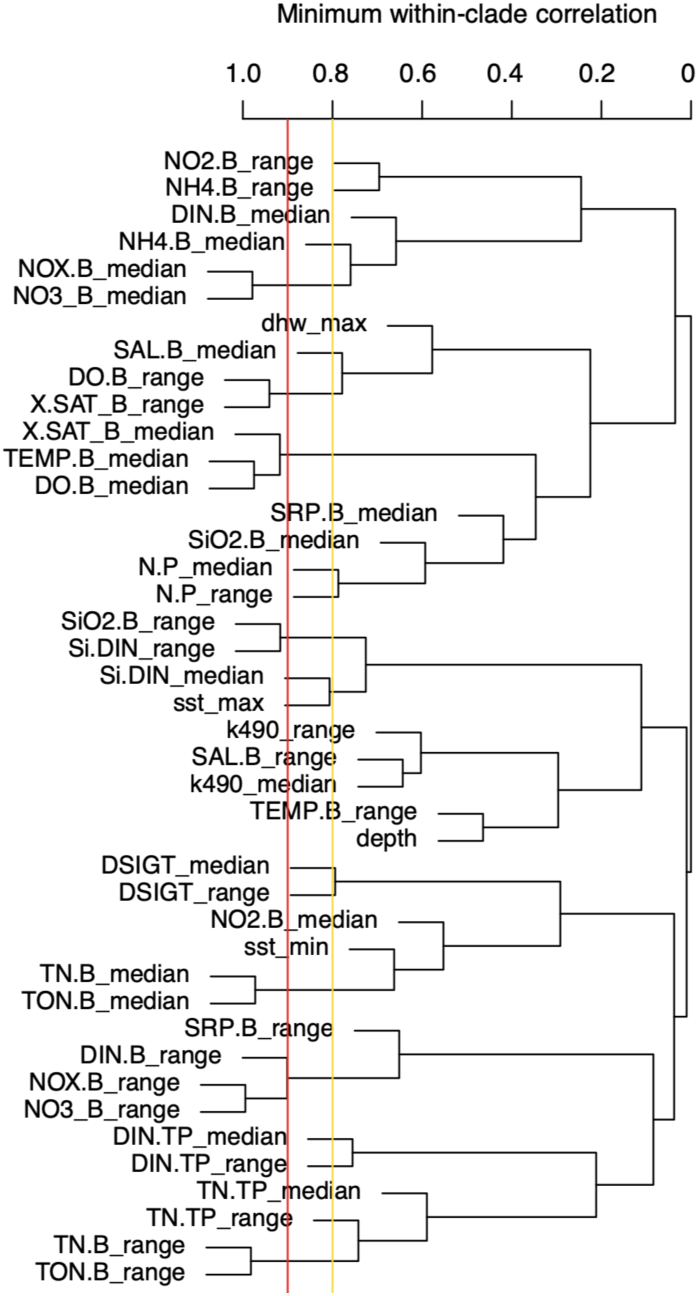
Hierarchical clustering tree of environmental variables. based on absolute correlation across sampled sites (complete linkage clustering). Variables joined into a clade to the left of the red line are correlated with Pearson’s *r* = 0.9 or higher.

**Supplementary Figure 3:**
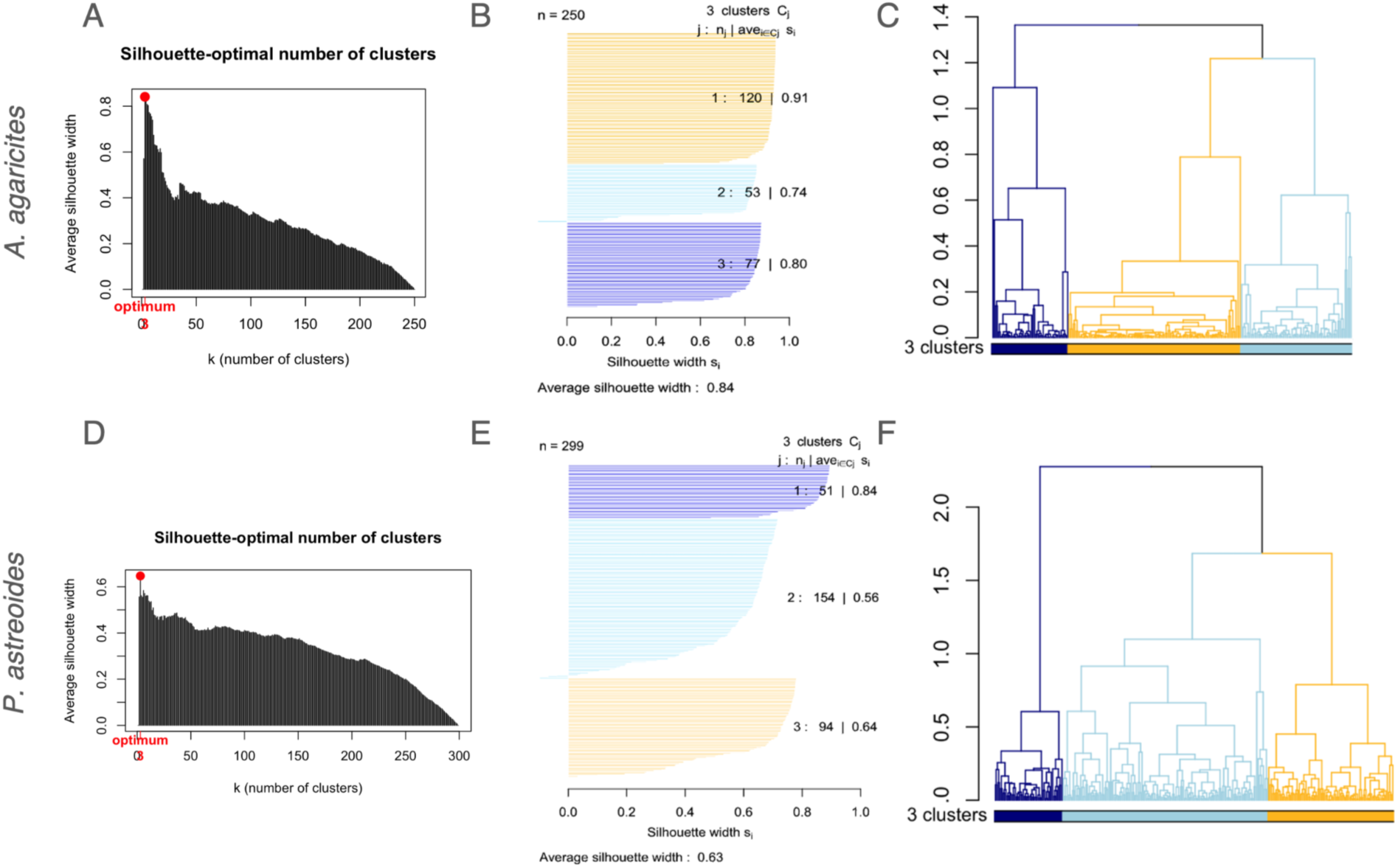
Optimal number of genetic clusters based on silhouette width (measure of the difference between within-cluster distances and distances to members of the next closest cluster). (A - C): *A. agaricites*, (D - F): *P. astreoides.* (A, D) Average silhouette widths for different numbers of clusters (*k*), based on the UPGMA clustering algorithm. In both cases, the widest silhouettes are observed at *k* = 3. (B, E) Silhouette widths for individual samples for *k* = 3. (C, F) Hierarchical clustering trees colored by cluster assignment.

**Supplementary Figure 4:**
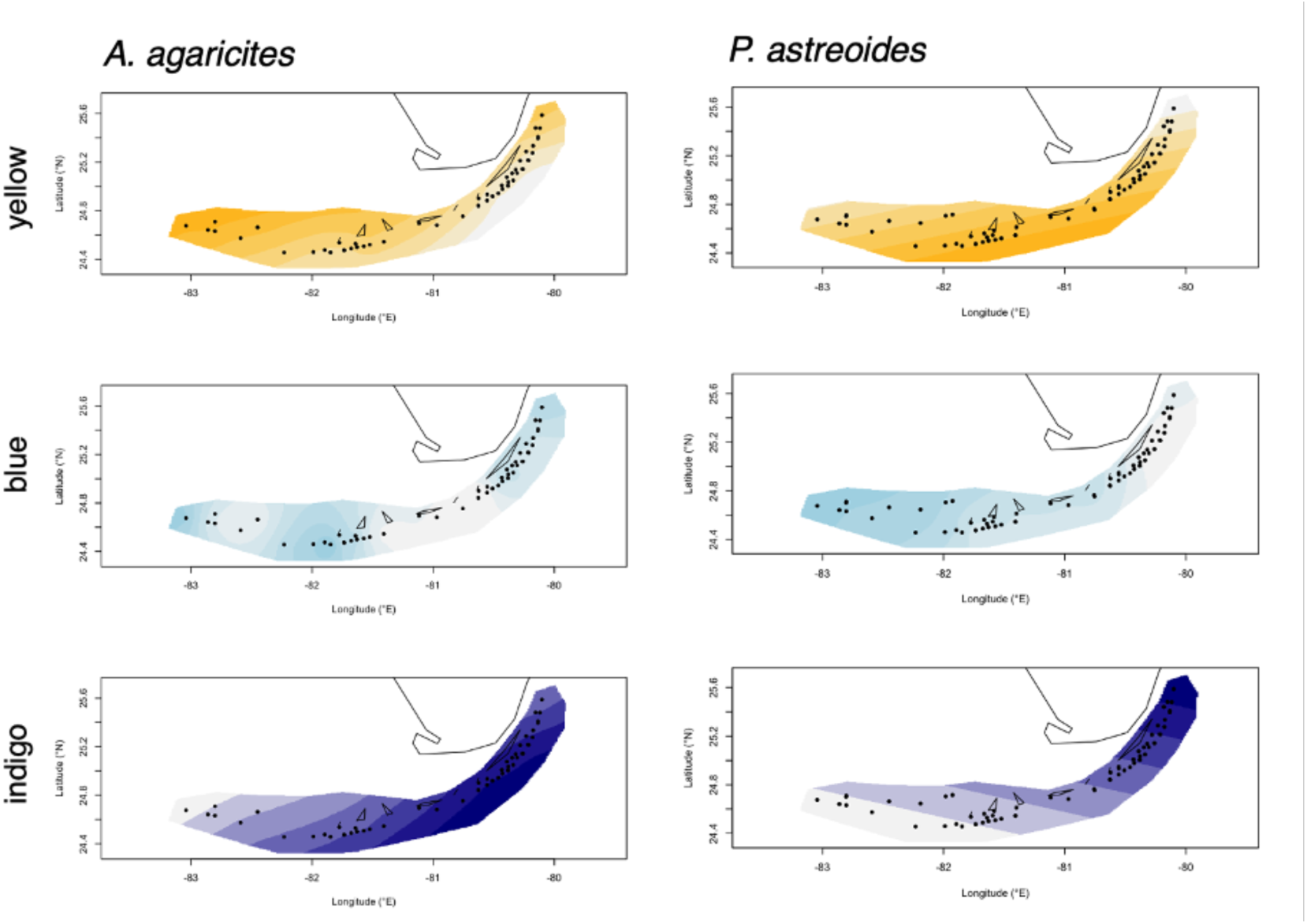
Distribution of genetic lineages of *A. agaricites* and *P. astreoides* across the FL Keys Reef Tract (FKRT). Spatial interpolation was performed using Tess3r (version 1.1.0, Caye et al., 2016) using ancestry proportions from Admixture analysis (Skotte et al., 2013). Dots indicate genetic sampling sites and more saturated colors indicate higher proportion of ancestry to each lineage. While distribution of lineage is uneven across the FKRT, they overlap broadly in their distribution.

**Supplementary Figure 5:**
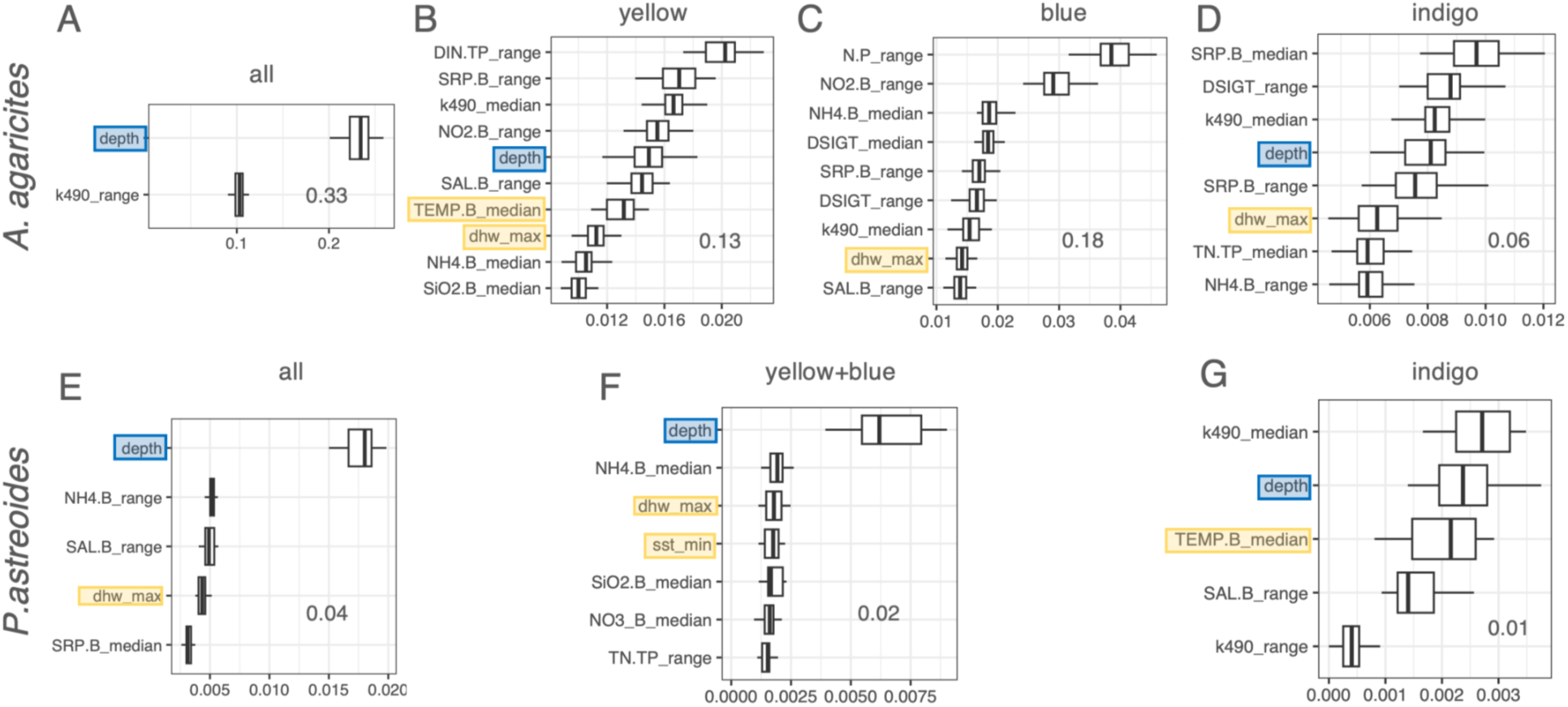
Proportion of total genetic variation explained by predictors retained after *mtry*-based selection procedure. (A-D) *Agaricia*, (E-G) *Porites*. The boxplots summarize importance (cross-validation *R*^2^) across 25 ordination jackknife replicates. Depth is labeled by blue rectangles, thermal variables – by yellow rectangles; all other variables describe water chemistry except k490 (turbidity) and DSIGT (water column stratification). Sample set is identified by the text label at the top of each panel: panels A and E show predictions based on all samples of a species, other panels show predictions based on subsets of samples corresponding to cryptic lineages.

**Supplementary Figure 6:**
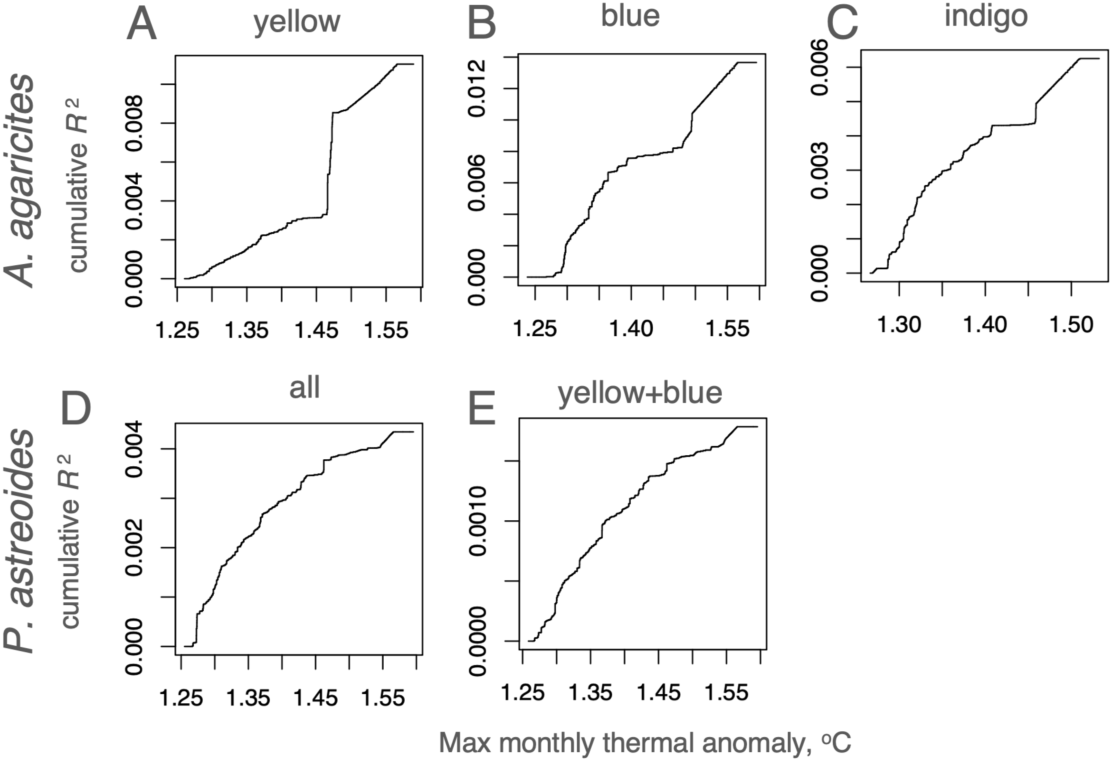
Turnover curves for maximum monthly thermal anomaly (dhw_max). (A-C) *Agaricia*, (D-E) *Porites*. Label on top of each panel identifies the sample set: either whole species (D) or cryptic lineages (A-C, E).

**Supplementary Figure 7.**
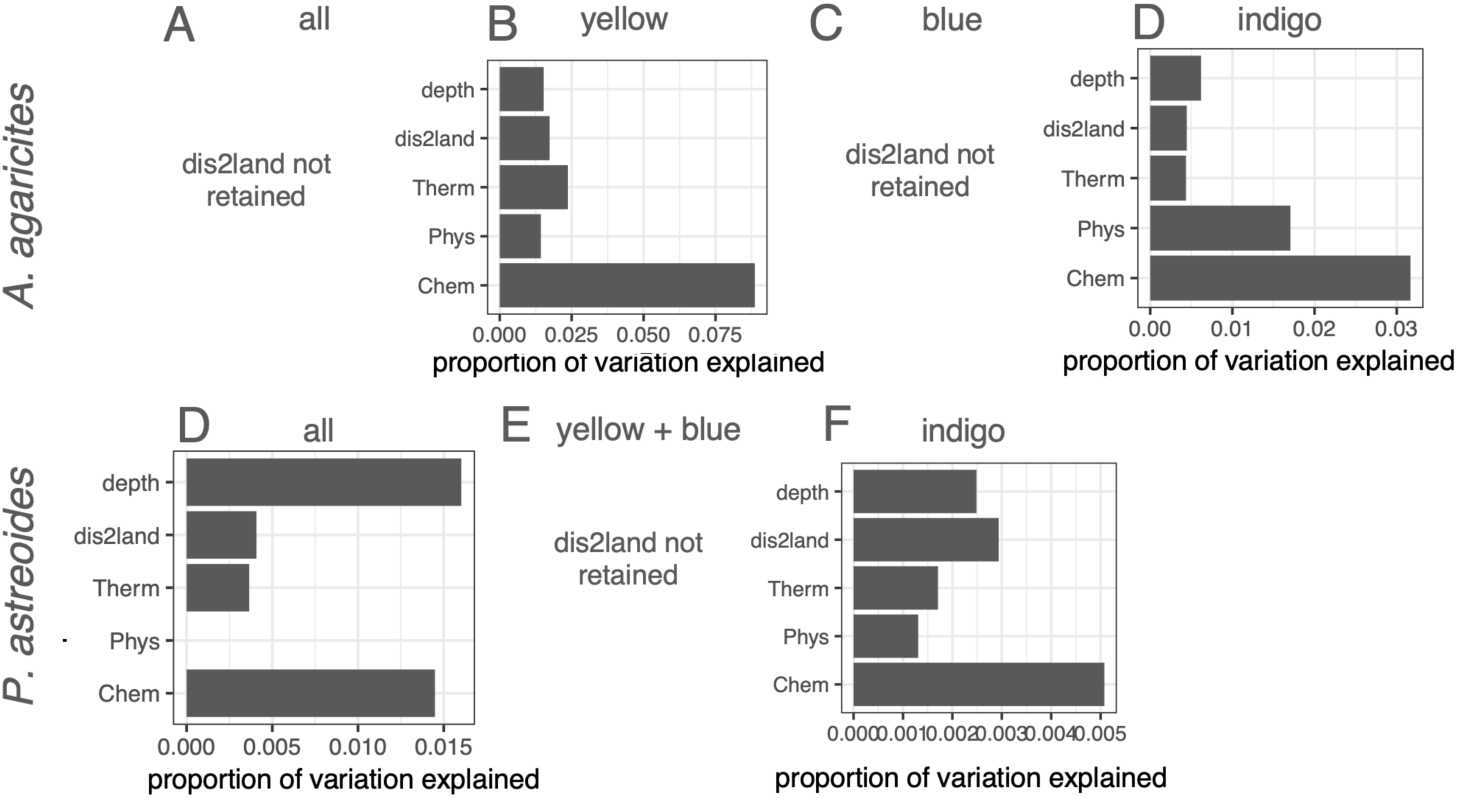
RDAforest models including distance to nearest land (dis2land) retain original predictors and do not substantially differ from original models. This Supplementary Figure hould be compared to Fig. 2. Bar charts are missing for analyses where dis2land did not pass the *mtry*-based selection criteria.

**Supplementary Figure 8:**
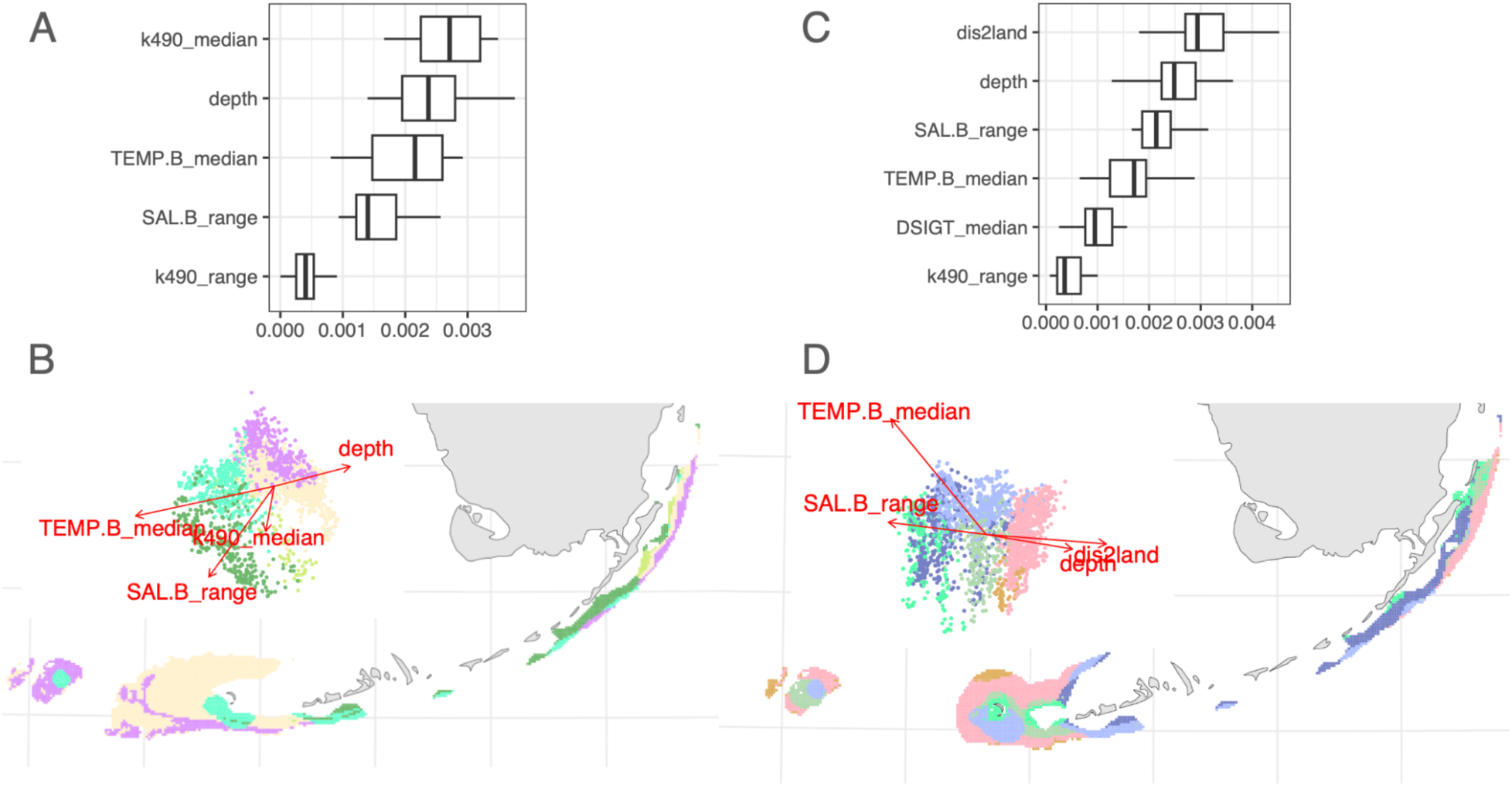
Distance to land does not invalidate specific environmental variables even if it is the most important predictor. Comparing importances of retained predictors (A, C) and maps of adaptive neighborhoods (B, D) for models for *P. astreoides* indigo either excluding distance to land (dis2land) from predictors (A, B), or including it (C, D).

**Supplementary Figure 9:**
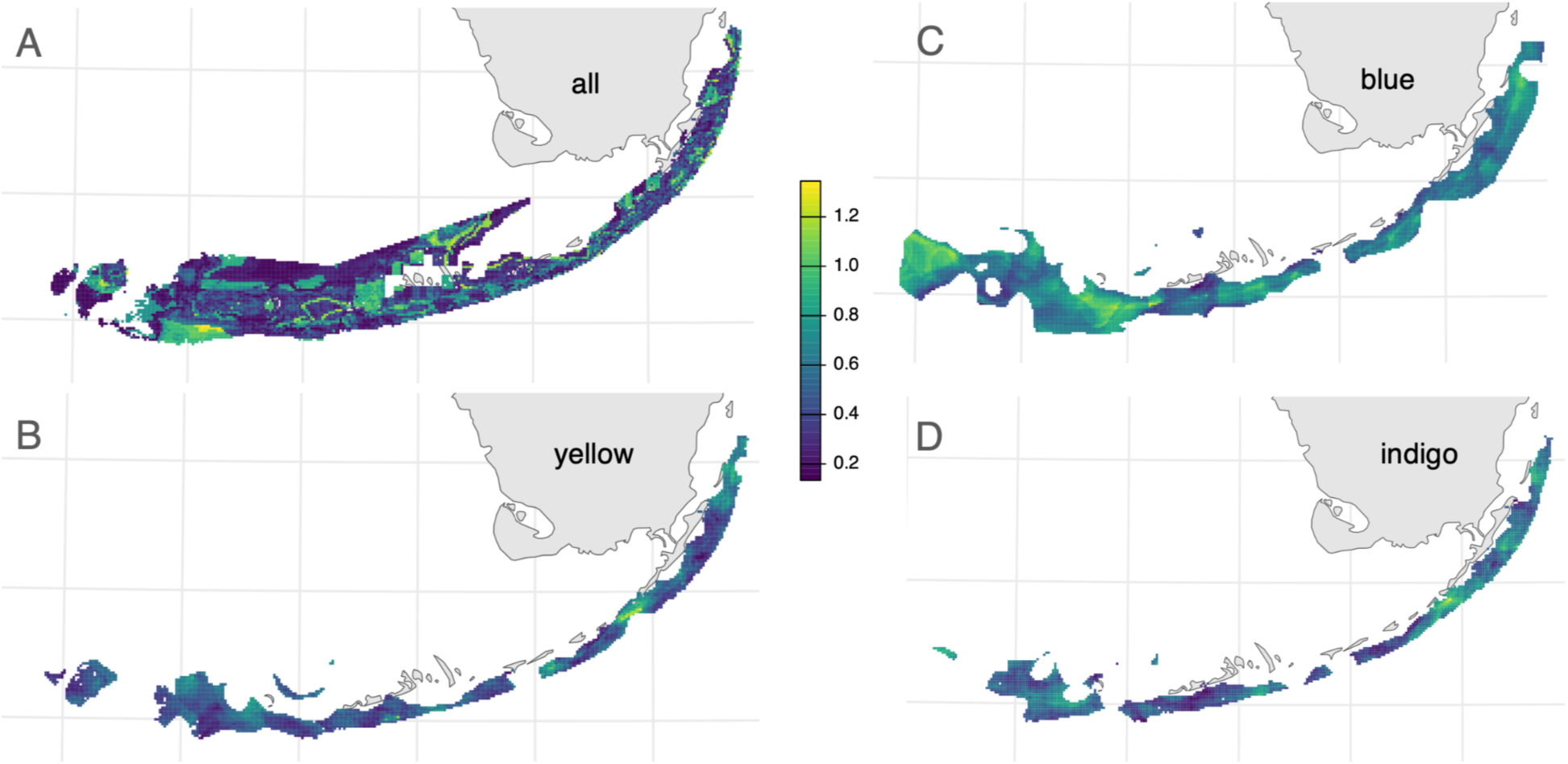
Genetic offsets for *A. agaricites*, for the whole dataset (A) and for individual cryptic lineages (B-D).

